# Leave-one-out cross-validation and linear modeling of visuospatial memory to predict long-term motor skill retention in individuals with and without chronic stroke: A short report

**DOI:** 10.1101/2020.10.14.330357

**Authors:** Jennapher Lingo VanGilder, Andrew Hooyman, Pamela R. Bosch, Sydney Y. Schaefer

## Abstract

Motor learning is fundamental to motor rehabilitation outcomes and has been associated with visuospatial memory function in previous studies. Current predictive models of motor recovery of individuals with stroke generally exclude cognitive measures, overlooking the connection between motor learning and visuospatial memory. Recent work has demonstrated that a clinical test of visuospatial memory (Rey-Osterrieth Complex Figure Delayed Recall) may predict one-month skill learning in older adults, but if this relationship persists in individuals with chronic stroke remains unknown. The purpose of this short report was to extend these findings by evaluating the extent these test scores impacted prediction in older adults and determine if this relationship generalized to individuals with stroke pathology. To address these questions, we trained two regression models (one including Delayed Recall scores and one without) using data from non-stroke older adults. To determine the extent to which Delayed Recall test scores impacted prediction accuracy of one-month skill learning in older adults, we used leave-one-out cross-validation to evaluate the prediction error between models. To determine if this predictive relationship persisted in individuals with chronic ischemic stroke, we then tested each trained model on an independent stroke dataset. Results indicated that in both stroke and non-stroke datasets, inclusion of Delayed Recall scores explained significantly more variance of one-month skill performance than models that included age, education, and baseline motor performance alone. This proof-of-concept suggests that the relationship between delayed visuospatial memory and one-month motor skill performance generalizes to individuals with chronic stroke and supports the idea that visuospatial testing may provide prognostic insight into motor rehabilitation outcomes.

## Introduction

Motor learning processes are fundamental to clinical motor rehabilitation. In other words, the benefits of motor therapy are theoretically predicated upon an individual’s capacity for skill reacquisition and long-term retention (1). Because the effects of stroke can vary greatly between individuals, responsiveness to motor therapy can be difficult to predict. There are already several models that have been developed to predict biological motor recovery post-stroke (e.g., the Predicting REcovery Potential algorithm (2)) that include personalized variables such as baseline motor function, age, severity of stroke, and white matter integrity. However, when attempting to predict changes in post-stroke upper-extremity impairment following therapy (i.e., responsiveness to motor therapy), recent work in machine learning has shown that the inclusion of sophisticated neuroimaging measures does not improve prediction accuracy beyond basic clinical measures (i.e., baseline Fugl-Meyer score) (3).

To our knowledge, no predictive models of therapeutic responsiveness include cognitive variables, despite growing evidence that they may explain significant amounts of variance in motor learning (4–6). For example, attention, executive function, and visuospatial memory underlie crucial stages of motor learning and are also among the most common cognitive deficits reported following stroke. Furthermore, a number of studies have shown that advancing age is associated with less improvement in motor therapy following stroke (7) and other musculoskeletal conditions. Since cognitive status often declines with age, it is plausible that responsiveness to motor therapy can, at least in part, be predicted by cognitive factors.

Empirically, there is a longstanding line of experimental motor learning studies that have shown that visuospatial function (i.e., of or relating to visual perception and spatial relationships between objects) is positively correlated with motor learning in both young and older adults (8–12). Our more recent work has begun to bridge the gap between empirical and clinical studies by showing that neuropsychological tests of visuospatial function may predict upper-extremity motor learning following task-specific training in older cohorts that are age-matched to a number of clinical stroke samples (e.g., (13)), whereas other clinical tests of attention, language, memory, etc., do not (14,15). This line of work has also highlighted that not all visuospatial tests are created equal, so to speak, since different visuospatial tests such as the Benton Judgement of Line Orientation (16) and the Wechsler Adult Intelligence Scale Block Design (17) probe different aspects of visuospatial function. By systematically comparing a battery of clinical visuospatial tests (including memory, perception, problem-solving, reasoning, and construction), we have demonstrated that only the Rey-Osterrieth Complex Figure Delayed Recall test (18), which measures visuospatial memory, uniquely predicted long-term skill retention of task-specific training in older adults without a history of stroke (13). This work strongly supports the premise that the same assessment (i.e., Delayed Recall) may also be a predictor of motor learning after stroke.

Thus, the purpose of this short report was to determine if the previously observed relationship between Delayed Recall test scores and one-month post-training skill performance in older adults persisted in individuals with a history of stroke and to evaluate the extent these test scores impacted prediction accuracy. This hypothesis-driven approach generated predictive models from a training dataset and then tested the generalizability of these models to an untrained dataset to test whether models that included visuospatial memory tests scores resulted in better predictive accuracy than models that did not.

## Methods

All experimental procedures were approved by Arizona State University’s Institutional Review Board and adhered to the Declaration of Helsinki. Forty-seven adults ages 56 to 87 years old (29 female/18 male) without a history of stroke comprised the training dataset, and seven adults with a history of ischemic stroke ages 33 to 81 (3 female/4 male) comprised the testing dataset. All participants provided informed consent prior study enrollment. A subset of data in the non-stroke cohort (n=45) has been published previously (13) and is included in the present study to model the predictive relationship described below. All participants were right-hand dominant (premorbidly if post-stroke), and were non-demented, based on established cut-off scores for neuropsychological assessments (see (13)). Participants with a history of ischemic stroke were also evaluated for motor deficits in their more-affected arm using the Upper Extremity Fugl-Meyer Assessment and the Action Research Arm Test. Post-stroke spasticity of the elbow flexors was evaluated using the Modified Ashworth Scale. Participants were excluded if they had hemispatial neglect, as determined by the Mesulam Cancellation Test. One participant had a right thalamic infarct, one had multifocal infarcts to the left middle cerebral artery related to high grade stenosis, one had a vertebral artery dissection, and one had a thrombotic ischemic stroke at the base of the cerebellum. Lesion location information was not available for three participants.

### Rey-Osterrieth Complex Figure Delayed Recall

This complex figure drawing test comprises two separate trials: A Figure Copy (measures visual construction) and a Delayed Recall (measures delayed visuospatial memory) trial. Participants were first asked to draw a replicate of a complex image as precisely as possible; once finished, all visual stimuli were removed from the testing area. Thirty minutes later, participants were asked to redraw the figure from memory (Fig. 1). To reduce interrater variability, a single rater scored each test using established testing guidelines. It is of note that the Delayed Recall score is independent of the Copy trial score (i.e., a high score on the Copy trial does not indicate the participant will achieve similar performance on the delayed memory trial). Based on our previous work using principal component analysis (13), only the Delayed Recall test scores were evaluated in this short report.

**Figure 1.**
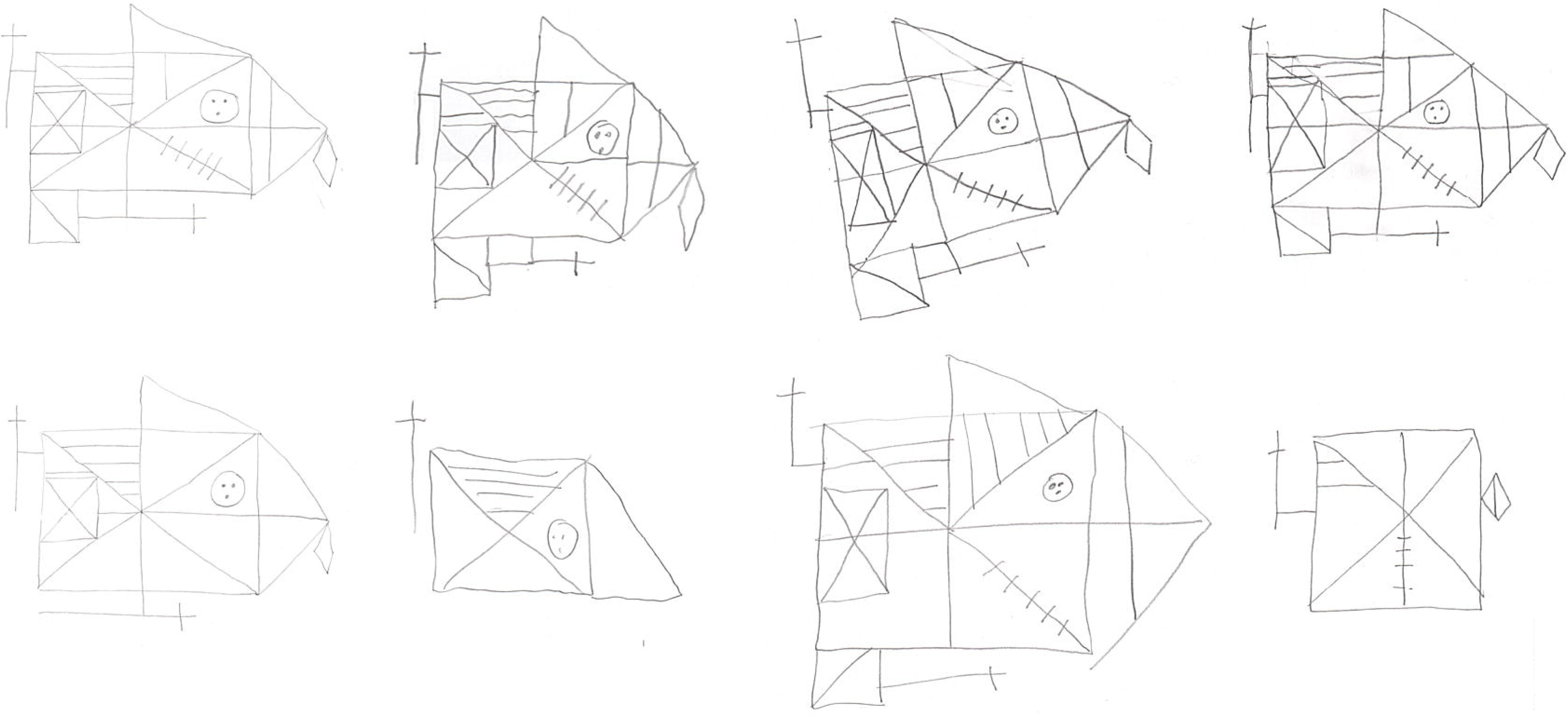
Participants completed the Rey-Osterrieth Complex Figure Copy (measures visual construction; on the top row) and Delayed Recall (measures visuospatial memory; on the bottom row); only the Delayed Recall trial was analyzed in this study. The Copy and Delayed Recall trials are scored independently from each other. Panels A and B show example drawings from older adults and individuals with a history of stroke, respectively. Note both groups demonstrated high performance on the Copy trial but marked variability in Delayed Recall performance.

### Task-specific motor training

Task-specific training included three sessions of 50 practice trials of a functional upper-extremity motor task over three consecutive weeks (one session/week). More details regarding the motor task are provided below. Participants were then re-tested one month after training to evaluate the amount of motor skill retained following a period of no practice; thus, our paradigm was designed with key principles of motor learning in mind, such as repetition and distributed practice. Furthermore, our paradigm is consistent with the goal of task-specific training, whereby participants practiced a functional task that simulated the basic activity of daily living of feeding oneself (19,20). To ensure the task was not overlearned, the non-stroke cohort used their nondominant hand; individuals in the stroke cohort used their more-affected hand. Given that all participants regardless of group were right-hand dominant, the non-stroke group performed all assessments and training with their left hand while most individuals in the stroke group did so with their right hand (one participant experienced left hemiplegia and used this hand accordingly).

The motor task used an experimental apparatus consisting of a wooden board (43 x 61 cm) with three different target cups placed radially around a constant ‘home’ cup at a distance of 16 cm (Fig. 2); each cup was 9.5 cm in diameter and 5.8 cm in height. Each trial began with thirty raw kidney beans in the home cup. The participant was instructed to pick up a standard plastic spoon located on the ipsilateral side of the home cup and use it to scoop two beans at a time from the home cup to the following sequence of target cups: ipsilateral, middle, then contralateral. This sequence was repeated until the last pair of beans were placed in the contralateral target cup, completing the trial. Errors such as transporting the wrong number of beans, dropping beans, or reaching in the wrong direction were recorded; error rates for both groups were modest (11.4% and 9.5% for stroke and non-stroke, respectively) and not included in our analyses. Participants were timed and instructed to move as quickly and as accurately as possible while freely exploring postural techniques to enhance performance (i.e., discovery learning). Trial time began when the participant picked up the spoon, with lower trial times indicating better performance. Since each trial consisted of 15 reaching movements, participants complete 750 reaches per training session, totaling 2,250 across the entire training paradigm. This task has ecological and construct validity (21) and instructional videos are available on Open Science Framework.

**Figure 2.**
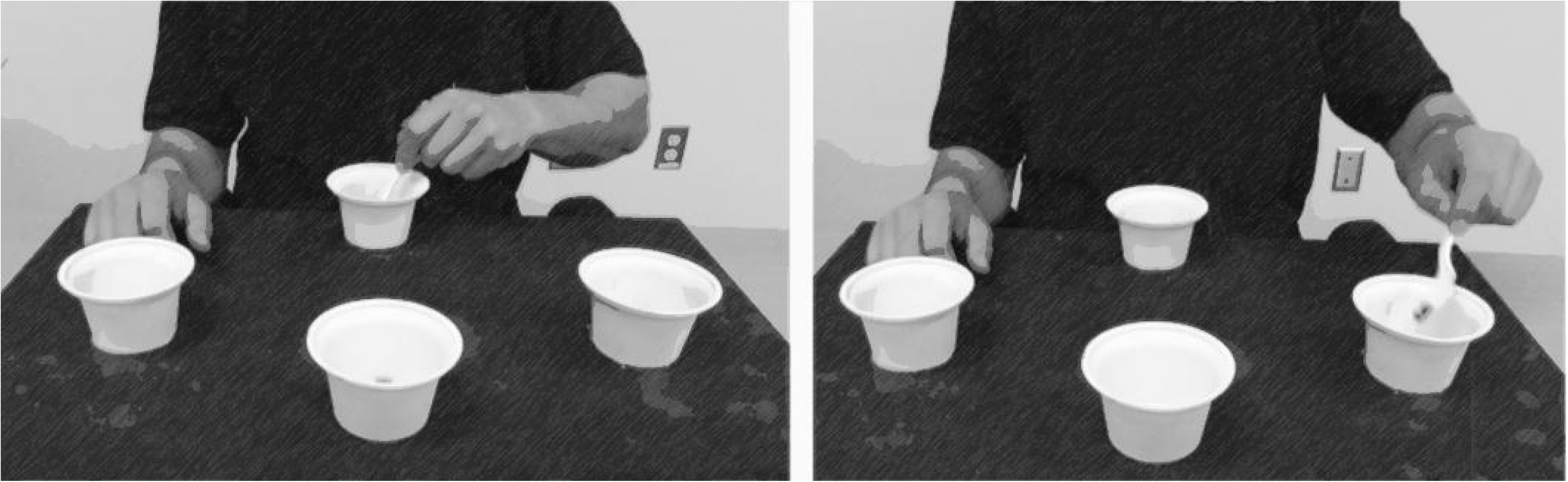
Participants used their nondominant hand to perform the motor task that mimicked the upper extremity movements required to feed oneself. This image is adapted from the “<u>Dexterity and Reaching Motor Tasks</u>” by <u>MRL Laboratory</u> that is licensed under <u>CC BY 2.0</u>.

### Statistical Analysis

Analyses were performed in JMP Pro 14.0 (SAS) and R Core Team 4.0.0 (2020) statistical software. To model the extent to which visuospatial memory test scores predicted one-month skill learning in a non-stroke cohort, multivariable regression was performed using covariates of age, education, Delayed Recall score, and baseline motor performance. Education was included to serve as a proxy for cognitive reserve (i.e., the brain’s resilience to neuropathological damage), which may explain differences in cognitive factors such as executive function, working memory, global cognition, and general arousal, as well as motor function following stroke (22). A separate model was then generated that excluded Delayed Recall scores to measure prediction accuracy without these visuospatial test scores, also in the non-stroke cohort. Analysis of variance (ANOVA) was then used to statistically compare prediction accuracy between both models. To test the robustness of this relationship, we performed two separate analyses: both models (Delayed Recall vs. no Delayed Recall) were 1) cross-validated in the non-stroke cohort using a leave-one-out approach (23) and 2) ‘trained’ using data from the non-stroke cohort and ‘tested’ on the independent stroke dataset. In leave-one-out crossvalidation, the model is trained on all data except that of a single participant and a prediction is made for that participant’s data; this process repeats for every participant (i.e., 47 times), thus all data are used for training the model but are used for prediction only once. This approach was chosen because it provides a method of generating unbiased prediction error to better estimate model fit. The mean squared error (MSE) between predicted versus observed values was calculated to compare accuracy among predictive models. This approach was designed to evaluate the extent to which visuospatial memory test scores can improve the prediction of longterm motor learning (i.e., comparison of MSE between Delayed Recall and no Delayed Recall models) and if this relationship generalizes to individuals with a history of stroke (i.e., comparison of MSE between non-stroke and stroke datasets).

The proposed approach has several strengths regarding rigor and reproducibility. First, by validating our model using data from our previous experiment (i.e., from a non-stroke cohort), bias is minimized. Second, by testing this validated model on an independent stroke dataset, the generalizability of this previously identified relationship can be examined within an independent clinical sample while minimizing the likelihood of statistical issues that are common in small sample sizes (e.g., a lack of statistical power, etc. (24)).

## Results

Participant characteristics, sensory and motor data are presented in Table 1. Overall, participants with a history of ischemic stroke had mild motor impairment, as indicated by their Upper Extremity Fugl-Meyer scores and their Action Research Arm Test scores. We acknowledge that the group had minimal motor deficits based on these stroke-specific assessments, but we point out, however, that participants with a history of ischemic stroke performed worse on the Grooved Pegboard Test than the non-stroke group (# drops, *p*=0.014; time to complete, *p*=0.093) even when performing it with their affected dominant (right) hand while the non-stroke group performed it with their nondominant (left) hand. Furthermore, the stroke group’s baseline performance on the motor task was worse than the non-stroke group’s performance with the same (right) hand (*p*=0.059). Collectively, these data indicate that the stroke group did in fact have some degree of motor impairment.

**Table 1.** Participant characteristics.

|  | *Control (nonstroke)* | *Stroke* |
| --- | --- | --- |
| Age<sup>a</sup> | 69.7±6.5<br>(56-87) | 58.4±16.5<br>(33-81) |
| Education<sup>b</sup> | 16.3±2.7<br>(12-24) | 15.9±2.0<br>(14-19) |
| Sex | 29 female<br>18 male | 3 female<br>4 male |
| Grip strength<sup>c</sup> | 29.1±11.1<br>(9.3-51.7) | 29.6±14.4<br>(11-56) |
| Grooved Pegboard Test<sup>d</sup> | 104.6±49.6<br>(64.6-335.6) | 138.7±48.8<br>(50.2-200.5) |
| Rey-Osterrieth Delayed Recall | 16.1±6.7<br>(2-31.5) | 12.3±8.3<br>(5-25) |
| Time post-stroke<sup>e</sup> |  | 3.8 ± 2.8<br>(1.1-9.7) |
| Upper extremity Fugl-Meyer<sup>f</sup> |  | 63 ± 3.5<br>(58-66) |
| Action Research Arm Test<sup>g</sup> |  | 52.7 ± 6.9<br>(38-57) |
| Modified Ashworth Scale<sup>h</sup> |  | 0.7 ± 1.1<br>(0-3) |

<sup>a</sup>in years; <sup>b</sup>in years; <sup>c</sup>in kilograms (dominant hand); <sup>d</sup>time, in seconds; <sup>e</sup>in years; <sup>f</sup>out of 66; <sup>g</sup>out of 57; <sup>h</sup>scale of 0-4, measuring elbow flexors

Motor training and retention data for participants with a history of stroke are presented in Figure 3 (it is noted that these data for non-stroke cohort have been published previously (13)). On average, participants improved performance from the baseline trial (mean±SD=53.09±11.31 seconds; 95% CI [44.72, 61.46]) to one-month follow-up (mean±SD=49.01±11.46 seconds; 95% CI [40.52, 57.50]), indicating that some participants improved more than others.

**Figure 3.**
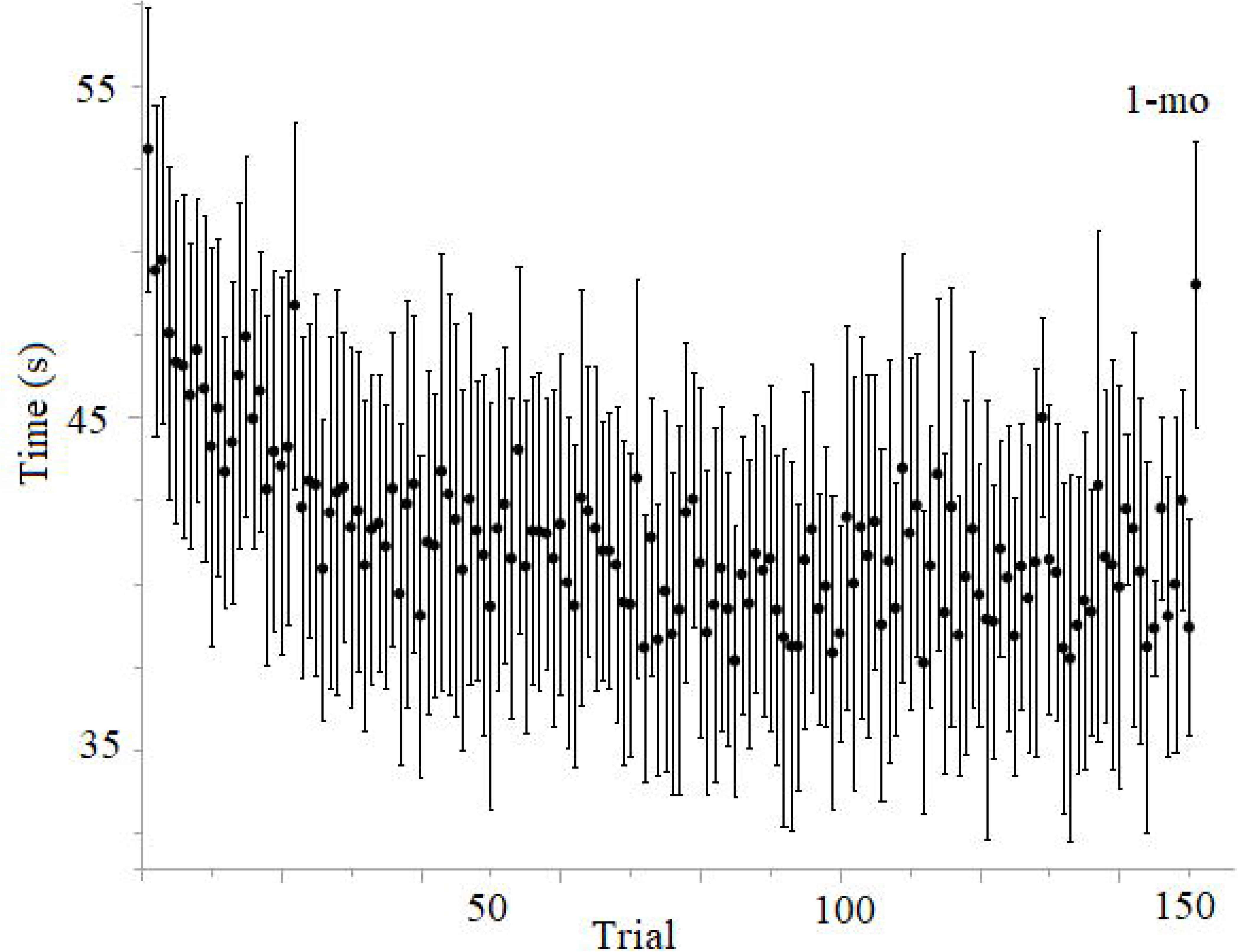
Participants completed a baseline trial of the reaching motor task, then completed 50 training trials during three weekly sessions (150 trials total). Participants were retested one month later to determine skill learning; four participants did not complete the third training session (trials 101-150) but did return for the one-month follow-up. Mean motor performance (trial time in seconds) is plotted on the y-axis, where lower values indicate better performance. Error bars indicate standard error.

To model the extent to which visuospatial memory predicted motor performance at one-month follow-up in the non-stroke cohort, multivariable regression included covariates of age, education, Delayed Recall score, and baseline motor performance (Table 2). Delayed Recall scores (*p*=0.025, □=-0.31; 95% CI [15.87, 16.38]) and baseline motor performance (*p*=0.002, □ = 0.31; 95% CI [58.12, 58.48]) demonstrated a similar effect size with one-month follow-up performance, where better scores predicted better performance at one-month follow-up. Age (*p*=0.22, □ = 0.19; 95% CI [-69.38, 70.01]) and education (*p*=0.67, □ = 0.13; 95% CI [-5.00, 16.98]) did not predict follow-up. In the comparison model that excluded Delayed Recall scores, only baseline performance (*p*<0.0001, □=0.36) predicted one-month follow-up performance (Table 3). ANOVA confirmed a significant difference between both models (*p*<0.05, Akaike information criterion of 305.4 vs. 308.4, respectively), indicating that the inclusion of Delayed Recall test scores explained more variance in motor performance at one-month follow-up than baseline, age and education alone, and improved the model’s overall goodness-of-fit.

**Table 2.** Parameters from the least-squares regression model including Delayed Recall that explain one-month follow-up performance.

| Based on 47 observations. |  |  |  |  |  |
| --- | --- | --- | --- | --- | --- |
| Parameter Estimates |  |  |  |  |  |
| *Name* | *Estimate ($\beta$)* | *SE* | *df* | *t-value* | *p-value* |
| Intercept | 19.47 | 12.39 | 0 | 1.57 | 0.124 |
| Delayed Recall | -0.31 | 0.13 | 1 | -2.33 | 0.025* |
| Age | 0.19 | 0.16 | 1 | 1.24 | 0.222 |
| Education | 0.13 | 0.34 | 1 | 0.43 | 0.673 |
| Baseline | 0.31 | 0.09 | 1 | 3.35 | 0.002* |

*Note.*

**Table 3.**
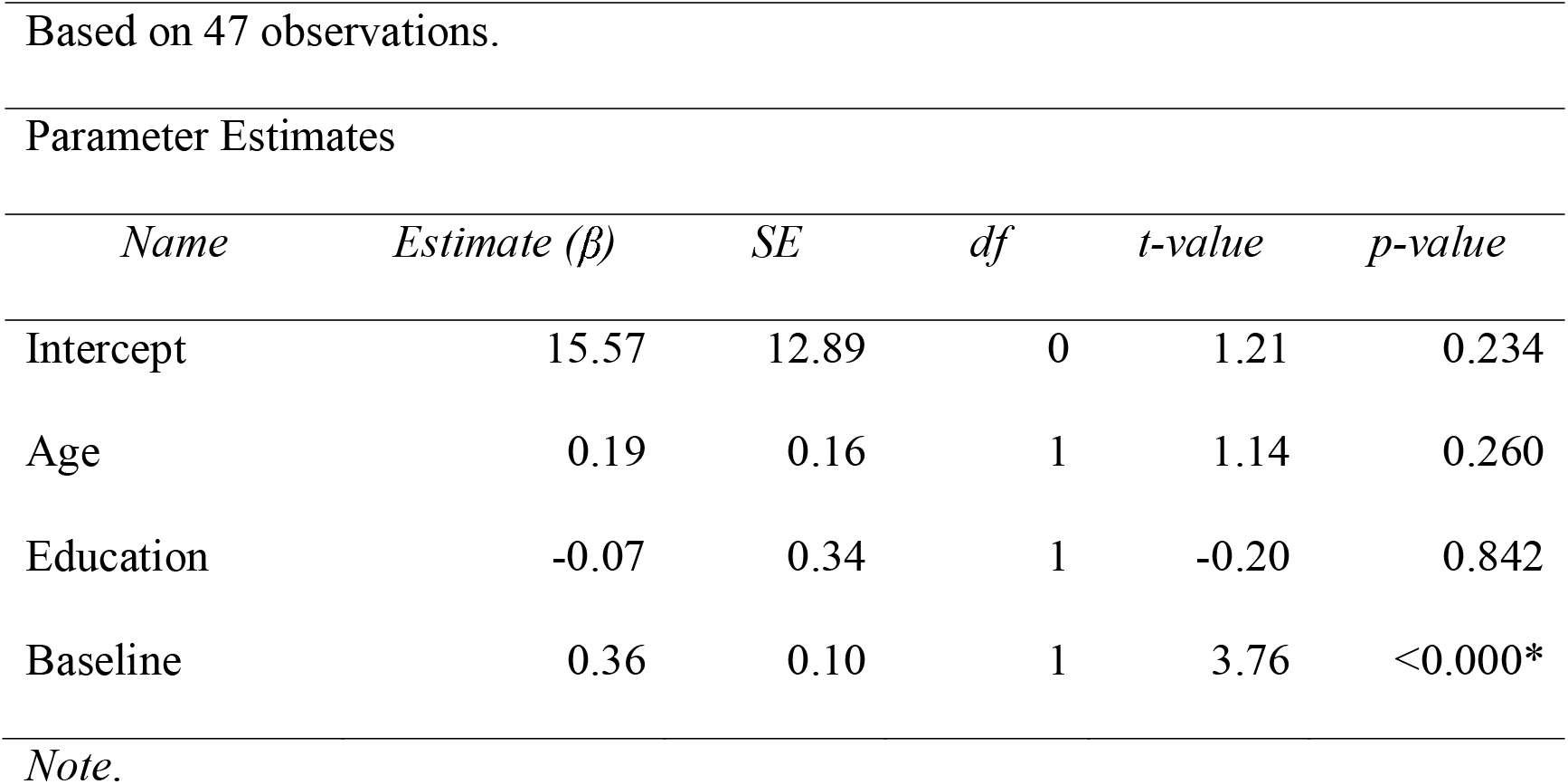
Parameters from the least-squares regression model excluding Delayed Recall that explain one-month follow-up performance.

To test the robustness of this relationship, both models (Delayed Recall vs. no Delayed Recall) were validated in the non-stroke cohort using a leave-one-out cross-validation approach. The mean squared error (MSE) between predicted and observed values for each model was 36.29 and 39.11 seconds, respectively (Fig. 4A). To test the generalizability of each model, both linear models (Delayed Recall vs. no Delayed Recall) were trained and then tested on the independent stroke dataset. The MSE between predicted and observed values for each model was 74.85 and 77.77 seconds, respectively (Fig. 4B). Overall, inclusion of Delayed Recall test scores reduced MSE, albeit modestly, in both models of one-month skill learning in non-stroke and stroke samples. However, these results from individual predictors are still of interest, given that the null model shows that group-based prediction performed much worse than individual-specific models. The resulting MSE was 87.24 seconds and 112.61 seconds for non-stroke and stroke groups, respectively.

**Figure 4.**
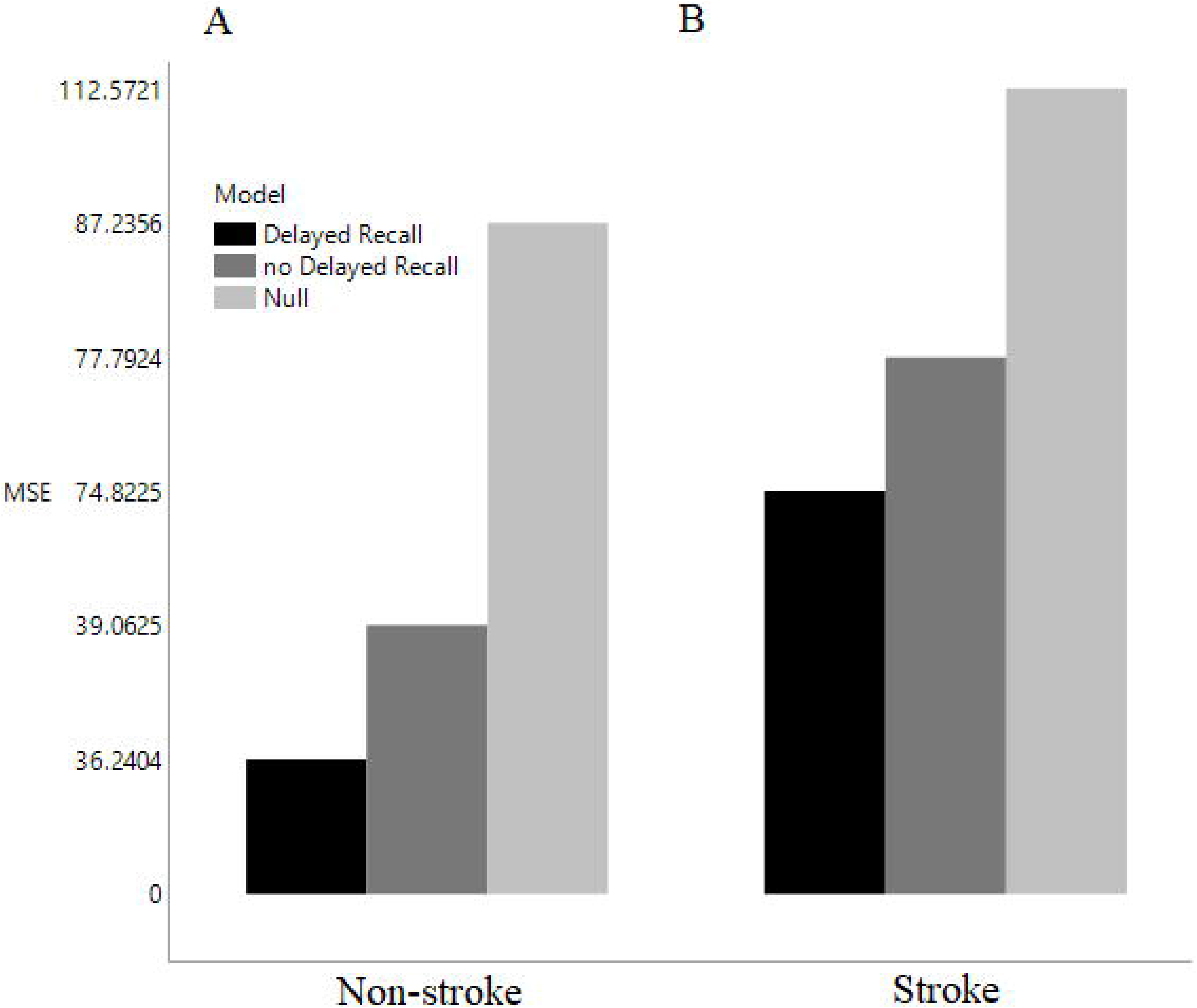
The mean squared error (MSE) for each model is presented for older adults and individuals with a history of stroke. A) Each model (Delayed Recall, no Delayed Recall, Null) was trained and tested on older adult data using a leave-one-out cross-validation approach; B) Each model was trained on older adult data and tested on individuals with stroke using a linear regression approach. Results indicate that in both groups, the inclusion of Delayed Recall test scores improved MSE.

## Discussion

The purpose of this short report was to determine the generalizability of the Rey-Osterrieth Complex Figure Delayed Recall test as a predictor of motor learning in a post-stroke cohort based on our previous findings, and to evaluate the extent to which adding these test scores as a predictor variable improved prediction accuracy. To address these hypothesis-driven questions, we trained two regression models (with and without Delayed Recall) using non-stroke data and tested them using leave-one-out cross-validation as well as on an independent stroke dataset using linear regression. Consistent with our hypothesis, results indicated that inclusion of Delayed Recall scores explained more variance in motor performance at one-month follow-up as compared to models that just included age, education, and baseline motor performance. This was consistent across both stroke and non-stroke datasets. These findings support the concept that visuospatial memory testing may provide prognostic insight into motor rehabilitation outcomes, and that cognitive rehabilitation could play a significant role in priming successful motor rehabilitation outcomes.

Despite the putative association between visuospatial memory and motor learning, cognitive variables are not currently considered in predictive models of upper-extremity motor recovery. This could be due to conflicting reports from clinical studies that evaluated the relationship between cognitive testing and motor rehabilitation outcomes. For example, change in motor outcomes has been linked to memory (25), executive (26), and visuospatial (27) functions, while other studies report no relationship between these cognitive domains and motor improvement (28). Comparison between reports is further confounded by differences in severity of impairment between groups, and more importantly, the lack of specificity in the cognitive tasks used. Often times global measures like the Montreal Cognitive Assessment or the MiniMental Status Exam are used to quantify cognition, but these tests insufficiently measure the function of specific cognitive domains especially pertinent to motor learning abilities and are often used as exclusion criteria (24).

A plausible mechanism underlying the association between visuospatial and motor learning is variation in structural integrity of specific white matter tracts among older adults and individuals with stroke. Structural neuroimaging studies in healthy aging (29) and stroke (30) demonstrate that white matter is particularly susceptible to the degenerative effects of normal aging and lesions. While the structural characteristics of frontoparietal white matter tracts have been linked to visuospatial function (31) and motor skill learning (32), it remains unknown if frontoparietal white matter microstructure explains variance in this behavioral relationship. Notably, the non-stroke group in this study used their left hand to complete the motor training, while the stroke group used their more affected hand. The fact that we observed a behavioral relationship between delayed visuospatial memory and one-month skill retention independent of which hand was used suggests that this effect is generalizable.

As with neural structures, motor learning and visuospatial function typically decline across the lifespan (33–37), yet one unexpected finding from the present study was that age did not demonstrate a significant effect on one-month follow-up performance. As a quality check, only participant age was included in the regression models of one-month follow-up, and results indicated that indeed age was related to follow-up performance (results not reported); we interpret this to suggest that behavioral factors such as baseline motor performance and delayed visuospatial memory are more sensitive predictors of one-month motor performance than chronological age (i.e., which explains why age is nonsignificant when these variables are included in the models). Moreover, our results indicate that visuospatial memory may explain variance beyond that of age, education, and baseline performance alone.

In regard to predicting spontaneous stroke recovery, this and other studies do not suggest that visuospatial memory scores can or should replace predictor variables used in current algorithms, or that the models presented here are valid recovery prediction tools; rather, the purpose of this short report was to demonstrate the predictive relationship between visuospatial memory and motor learning persists in individuals with a history of stroke, and to empirically support the premise that visuospatial memory testing may be an overlooked consideration for understanding why responsiveness to motor rehabilitation can be so varied.

Previous research has shown that effect sizes and beta values derived from a small sample group are highly prone to inflation and therefore may be unreliable (24). To avoid this pitfall in our analyses, we first evaluated the reliability of the behavioral relationship in a moderately large non-stroke group using leave-one-out cross validation; results indicated beta values in this dataset were reliable. This validated model was then used to test if the behavioral relationship also generalizes to individuals with a history of stroke. In other words, while the stroke cohort in this study was small, our analyses were not wholly dependent upon its sample size and the potential limitations associated with it. Another limitation to this study is that all participants were in the chronic stage of stroke (>1 year) and exhibited very mild motor impairment. It is possible that individuals with more moderate-to-severe motor impairment (had we been able to recruit them prior to the COVID-19 shutdown) would have also had more impaired visuospatial ability (38), which would support previous findings of less motor learning with higher stroke severity (39) based on our working hypothesis and regression model. However, as noted above, a larger, more acute, and more impaired sample has not been recruited due to COVID-19, preventing us from directly testing whether this model would retain comparable prediction accuracy in more acute or more impaired individuals. This is not a trivial question, since different cognitive deficits tend to emerge throughout the recovery process (e.g., attention deficits during acute (40) and visuospatial and memory deficits present at three months post-stroke (41)). Thus, the current study cannot discern if the presence of other cognitive impairments will impact this behavioral relationship. However, since the Rey-Osterrieth Complex Figure Delayed Recall test is a validated measure of nonverbal memory, executive function, and graphomotor skills, it likely captures the breadth of cognitive impairments most common following stroke. To address these limitations, future work will involve recruiting a larger and more impaired sample and evaluating if Delayed Recall scores, and other specific neuropsychological tests, can be used to improve prediction in models involving individuals with stroke during the acute stage.

### Conclusions

In summary, the inclusion of Delayed Recall test scores modestly improved the accuracy in predictive models of one-month skill learning in individuals with and without stroke. These findings support the concept that visuospatial memory testing may provide prognostic insight into upper extremity motor learning and encourage future work to examine the role of cognitive testing in predictive models of motor recovery.

## Declarations

### Ethics approval and consent to participate

The participant recruitment and experimental protocols for this study was approved by Arizona State University’s Institutional Review Board. All protocols aligned with the Declaration of Helsinki and participants provided informed consent prior study enrollment.

### Consent for publication

Not applicable.

### Availability of data and materials

The datasets used and/or analyzed during the current study are available from the corresponding author on reasonable request.

### Authors’ contributions

JLV conceptualized the project, analyzed and interpreted data, and wrote the original draft. AH analyzed and interpreted the data and read and approved the final manuscript. PRB assisted in participant recruitment and clinical assessment. SYS supported the project, interpreted data and reviewed and edited the final draft. All authors read and approved the final manuscript.

## List of abbreviations

ANOVA: analysis of variance
MSE: mean squared error

